## Supplementary figures and images for "Appropriate glycemic management protects the germline but not uterine environment in type 1 diabetes"

### Figure S1

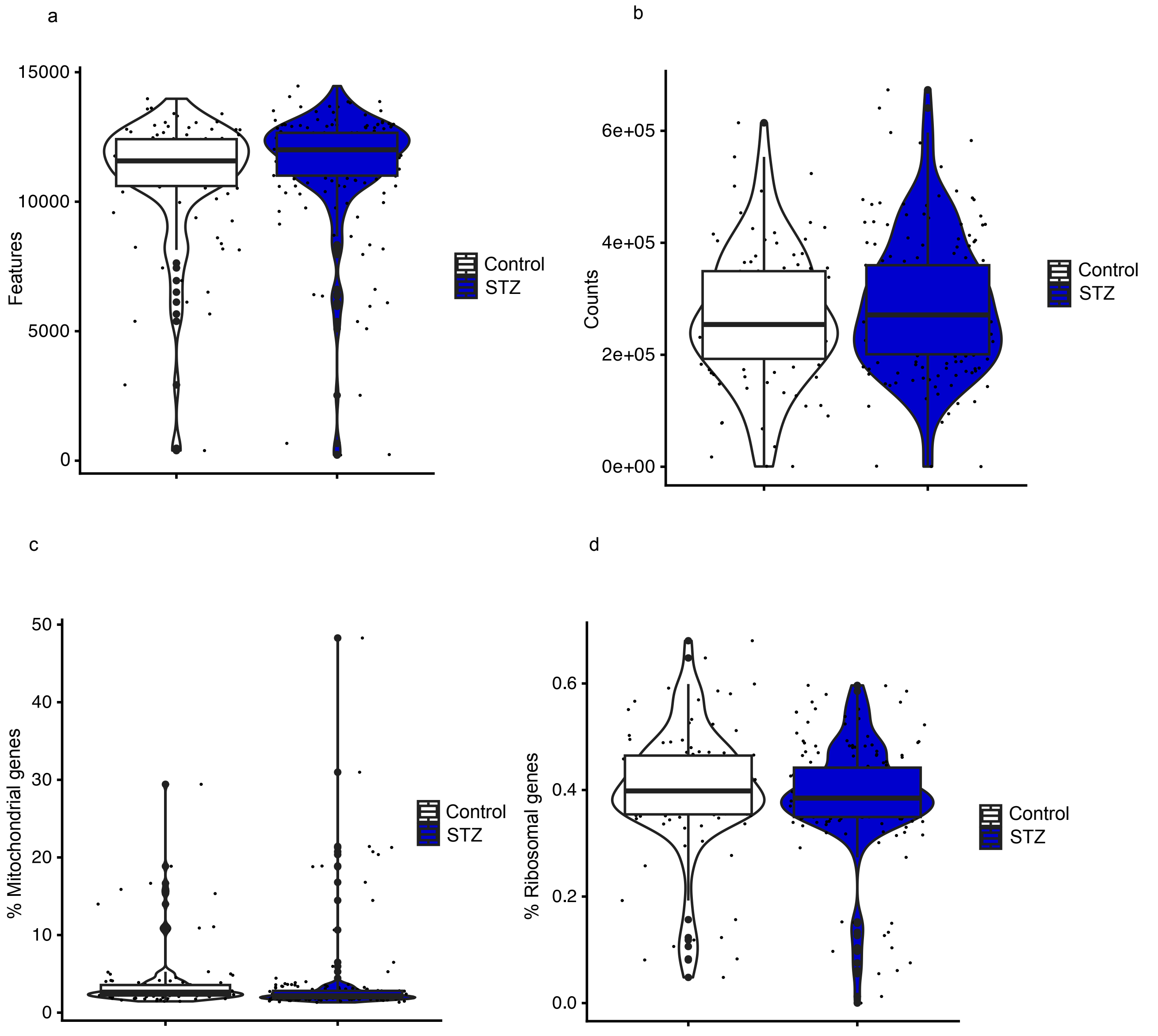

### Figure S2

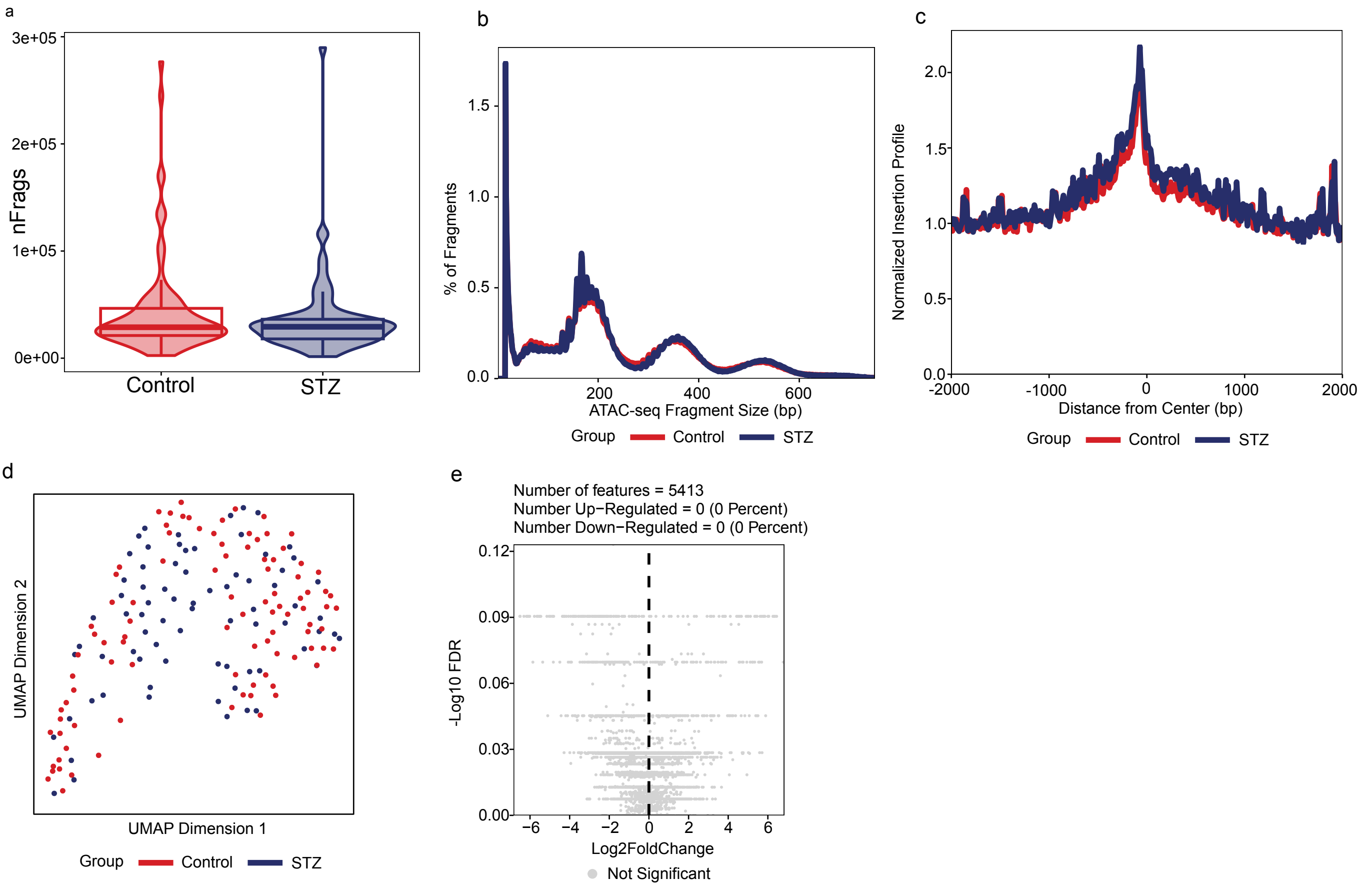

### Figure S3

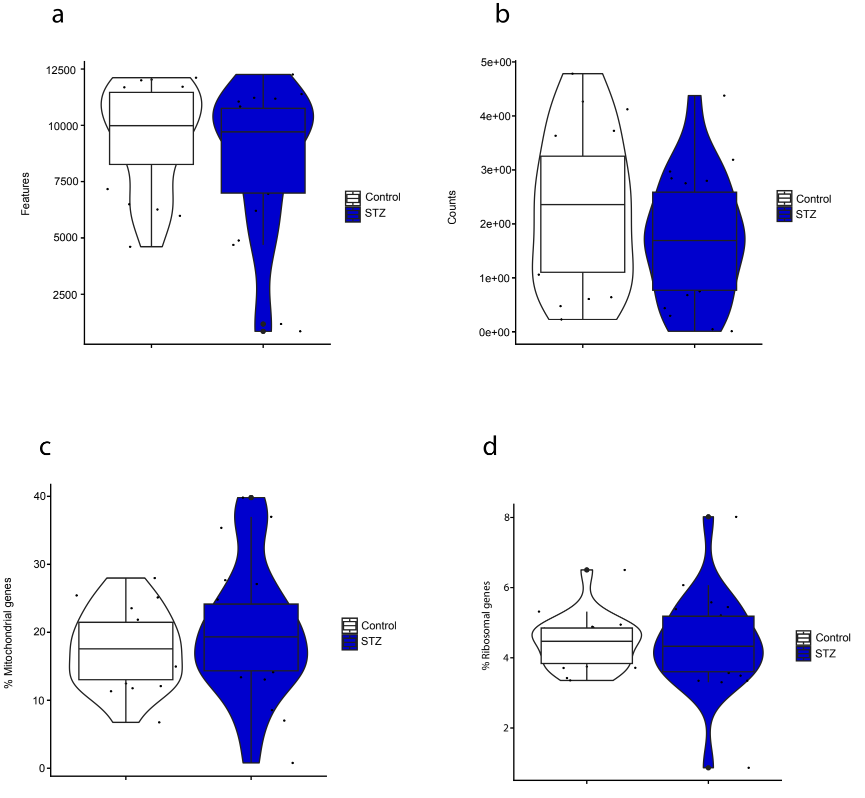
